# AAV-mediated neuronal expression of FOXG1 restores oligodendrocyte maturation, myelination, and hippocampal structure in mouse models of FOXG1 syndrome

**DOI:** 10.64898/2025.12.04.692422

**Authors:** Jaein Park, Holly O’Shea, Shin Jeon, Dongjun Shin, Liwen Li, Seon Ung Hwang, Michael Kofi Anyane-Yeboa, Songlin Yang, Camille F. Harrison, Yeong Shin Yim, Jae W. Lee, Soo-Kyung Lee

## Abstract

FOXG1 syndrome is a devastating neurodevelopmental disorder caused by haploinsufficiency of the transcription factor FOXG1, leading to intellectual disability, epilepsy, and white-matter deficits. Although FOXG1 is well known for its neuronal functions, its role in glial pathology remains poorly understood. Here, we show that reducing FOXG1 selectively in neurons impairs oligodendrocyte lineage progression and myelination, establishing a critical non-cell-autonomous role for neuronal FOXG1 in glial maturation. To restore FOXG1 in neurons, we developed AAV vectors expressing human *FOXG1* under neuron-specific promoters. Neonatal administration of these vectors normalized oligodendrocyte precursor cell (OPC) accumulation, enhanced myelination, and corrected hippocampal structural abnormalities in *Foxg1* conditional heterozygous mice. To test therapeutic robustness under stringent conditions, we used the patient-specific W300X heterozygous model, which combines FOXG1 loss-of-function with a toxic truncated protein and represents one of the most severe FOXG1 syndrome genotypes. Remarkably, neuron-restricted AAV-FOXG1 delivery produced substantial rescue even in this high-bar model, suppressing OPC overaccumulation, restoring myelination, and progressively improving dentate gyrus morphology, with benefits persisting into adulthood. Moreover, adolescent administration remained highly effective, rescuing myelination, axonal bundle thickness, and microglial activation. These findings identify neuronal FOXG1 as a master regulator of neuron-glia interactions and establish neuron-targeted AAV-FOXG1 as a potent and clinically translatable therapeutic strategy across diverse severities of FOXG1 syndrome.

## INTRODUCTION

The transcription factor FOXG1 is a pivotal regulator of forebrain development, controlling the balance between neural progenitor cell (NPC) self-renewal and neuronal differentiation. In NPCs, FOXG1 regulates proliferation and the timing of neurogenesis, thereby influencing the size and structure of the cerebral cortex (Xuan et al. 1995; Hanashima et al. 2002; Martynoga et al. 2005; Manuel et al. 2011; Hettige et al. 2022). In postmitotic neurons, FOXG1 orchestrates neuronal migration, axonal projection, dendritic maturation, and synaptic development (Miyoshi and Fishell 2012; Kumamoto et al. 2013; Cargnin et al. 2018; Shen et al. 2019; Ni et al. 2021). Additionally, it promotes neuronal survival and activity-dependent plasticity (Dastidar et al. 2011; Dastidar et al. 2012; Yu et al. 2019; Tigani et al. 2020).

Within the hippocampus, FOXG1 exerts multiple functions essential for both development and lifelong plasticity. It plays critical roles in hippocampal morphogenesis and patterning, regulating the expansion and fate of neural stem cells (NSCs) in the dentate gyrus (DG), a region vital for adult neurogenesis, learning, and memory (Manuel et al. 2010; Tian et al. 2012). FOXG1 further influences the maturation of postmitotic hippocampal neurons, coordinating dendritic arborization, axon projection, and synaptic connectivity (Ni et al. 2021; Akol et al. 2023). In the adult hippocampus, FOXG1 continues to support neuronal survival, neurogenesis, and synaptic plasticity, underscoring its enduring importance for hippocampal structure and function (Yu et al. 2019; Wang et al. 2022a; Schäffner et al. 2023).

While the cell-autonomous functions of FOXG1 in neuronal differentiation are well established, its influence on glial biology remains poorly understood. FOXG1 has been shown to inhibit astrocyte differentiation from neural progenitor cells (Brancaccio et al. 2010; Falcone et al. 2019; Bose et al. 2025), but its role in the oligodendrocyte lineage has not been defined. This represents an important knowledge gap, as oligodendrocyte development and myelination are essential for neuronal function and network integrity. Addressing this question is also clinically significant, given that FOXG1 haploinsufficiency in humans is consistently associated with myelination deficits (Harada et al. 2018; Mitter et al. 2018; Vegas et al. 2018; Pringsheim et al. 2019).

Emerging evidence suggests that FOXG1-driven processes are expression level dependent, particularly in the hippocampus and DG. Conditional heterozygous deletion of *Foxg1* in cortical and hippocampal neurons results in defects in DG morphogenesis and function (Cargnin et al. 2018; Jeon et al. 2024). Global heterozygous *Foxg1* mutants (*Foxg1*^+/−^) exhibit microcephaly, reduced cortical thickness, hippocampal abnormalities, cortical and hippocampal hyperexcitability, and behavioral deficits (Shen et al. 2006; Eagleson et al. 2007; Siegenthaler et al. 2008; Pringsheim et al. 2019; Testa et al. 2019a; Testa et al. 2019b; Miyoshi et al. 2021; Erickson et al. 2022; Younger et al. 2022). More recently, patient-specific models carrying FOXG1 syndrome mutations highlighted the importance of FOXG1 expression levels in the oligodendrocyte lineage: Q84Pfs heterozygous mice show aberrant accumulation of oligodendrocyte precursor cells (OPCs) accompanied by reduced myelination (Jeon et al. 2025b). These findings point to an essential role of FOXG1 in regulating oligodendrocyte lineage progression and suggest that hypomyelination may arise from impaired OPC differentiation.

These experimental insights are directly relevant to FOXG1 syndrome, a severe neurodevelopmental disorder caused by heterozygous pathogenic variants in *FOXG1*. Patients present with profound intellectual disability, early-onset epilepsy, and structural brain anomalies including microcephaly and corpus callosum hypogenesis (Kortum et al. 2011). Neuroimaging consistently reveals hippocampal malformations and white-matter reduction (Harada et al. 2018; Mitter et al. 2018; Vegas et al. 2018; Pringsheim et al. 2019). Importantly, myelination deficits are a constant feature in patients with FOXG1 syndrome. In two fetal cases, neuropathological examination showed an increase in OPCs with a corresponding reduction in pre-myelinating oligodendrocytes (Wilpert et al. 2021), consistent with a differentiation block. Together, human and mouse studies suggest that FOXG1 haploinsufficiency impairs oligodendrocyte maturation and myelin development, adding glial dysfunction to the well-recognized neuronal pathology in FOXG1 syndrome.

Here, we address this gap by investigating whether neuronal FOXG1 expression alone can restore both neuronal and glial functions in FOXG1-deficient brains. Using adeno-associated virus (AAV) vectors engineered to express human *FOXG1* under neuron-specific promoters, we demonstrate that neuronal FOXG1 restoration is sufficient to correct widespread pathology across multiple FOXG1 syndrome models. In conditional *Foxg1* heterozygous mice, neuron-restricted AAV-FOXG1 normalized oligodendrocyte lineage progression, enhanced myelination, and rescued hippocampal architecture with efficacy comparable to AAV-FOXG1 driven by the ubiquitous CBA promoter (Jeon et al. 2024). Notably, the same neuron-specific strategy proved equally effective in the patient-derived W300X model, a stringent allele combining FOXG1 loss-and gain-of-function mechanisms, fully restoring myelination and oligodendrocyte differentiation, even when administered during adolescence. These results establish neuronal FOXG1 as a principal regulator of neuron-glia interactions and demonstrate that targeting neurons alone is sufficient for broad therapeutic benefit. Together, these findings provide compelling preclinical evidence that neuron-targeted AAV-FOXG1 gene replacement is a potent and clinically translatable therapeutic approach for FOXG1 syndrome.

## RESULTS

### Generation of AAV-FOXG1 vectors that drive human *FOXG1* expression in neurons

To test whether FOXG1 syndrome phenotypes can be ameliorated by restoring FOXG1 postnatally, we generated new AAV vectors encoding codon-optimized human *FOXG1* cDNA. Given that the neuronal action of FOXG1 expression is a key determinant of forebrain development, we designed two configurations incorporating neuron-selective promoters: Synapsin1 promoter (pSYN1) (Kügler et al. 2003; Boulos et al. 2006), and a modified chicken β-actin (CBA) promoter harboring tandem repressor element 1 (RE1) sites, which restrict promoter activity to neurons by suppressing transcription in non-neuronal cells (Bessis et al. 1997; Millecamps et al. 1999) (Fig. 1A).

**Figure 1.**
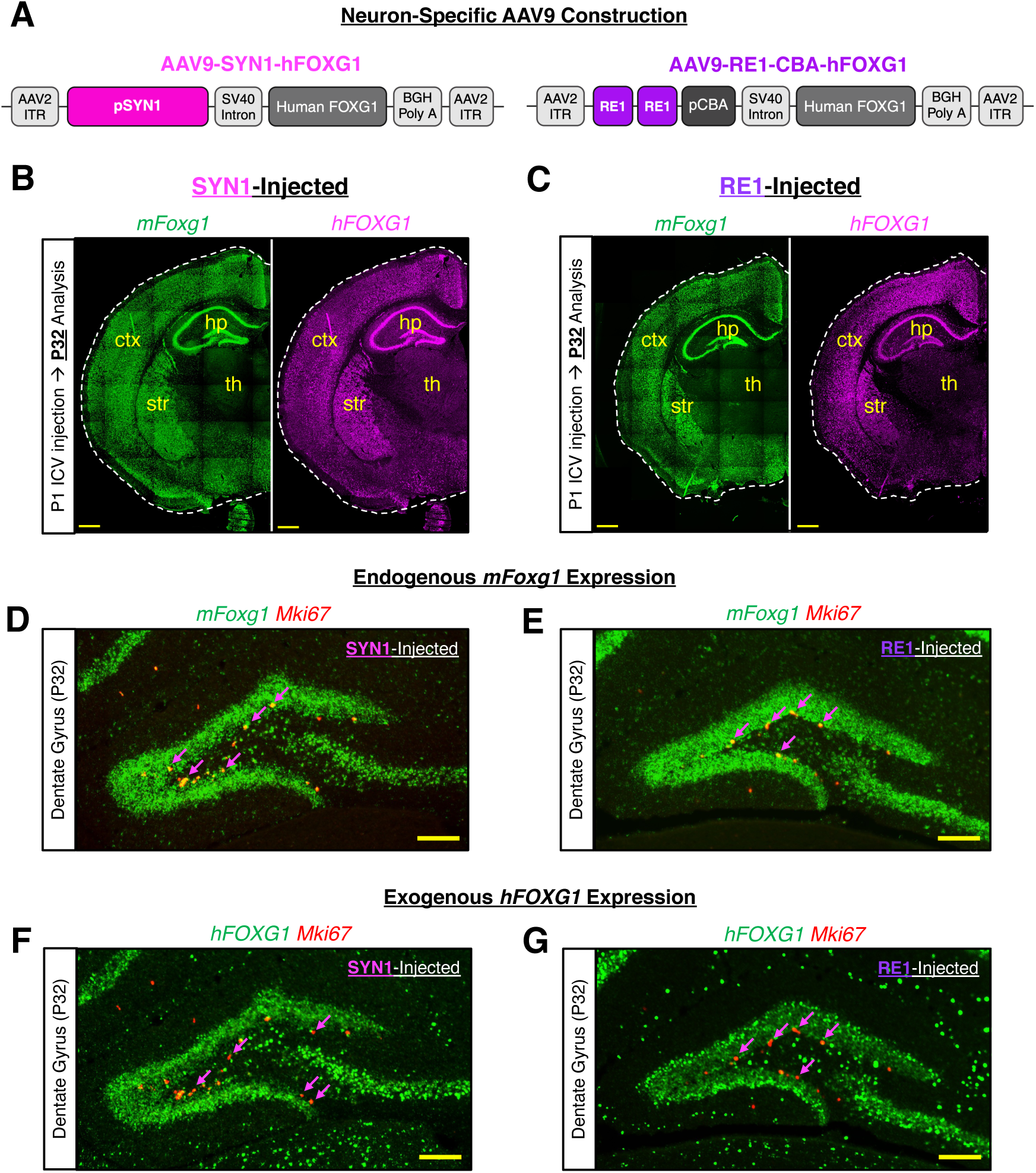
AAV-SYN1 and AAV-RE1 successfully drive neuron-specific expression of human *FOXG1 in vivo* in control brains. (A) Schematic representations of SYN1 and RE1 vector designs. (B-C) *In situ* hybridization analysis comparing endogenous mouse *Foxg1* expression with virus-driven human *FOXG1* expression shows that P1 ICV injection of SYN1 or RE1 drives *hFOXG1* expression in biologically relevant brain areas at P32. Scale bar 500µm. ctx, cortex; hp, hippocampus; str, striatum; th, thalamus. (D-E) *In situ* hybridization analysis detects endogenous *mFoxg1* expression overlapping with proliferation marker *Mki67* in the P32 dentate gyrus. Arrows indicate *mFoxg1^+^/Mki67^+^*cells. Scale bar 200µm. (F-G) *In situ* hybridization analysis shows that SYN1 and RE1-driven *hFOXG1* is not co-expressed with *Mki67^+^*cells in the P32 dentate gyrus, indicating neuronal selectivity. Arrows indicate *hFOXG1^-^/Mki67^+^* cells. Scale bar 200µm. n = 3 mice per condition. Representative images shown.

To assess if AAV-pSYN1-FOXG1 (hereafter termed SYN1) and AAV-RE1-pCBA-FOXG1 (termed RE1) viruses drive exogenous human FOXG1 expression in neurons, we performed intracerebroventricular (ICV) injection of SYN1 or RE1 virus in control newborn mice. We then analyzed the expression pattern of human *FOXG1* in P32 brains using double immunofluorescence *in situ* hybridization (ISH) assays (Fig. 1B,C). We used human *FOXG1-* and mouse *Foxg1*-specific probes that detect exogenous human *FOXG1* and endogenous mouse *Foxg1* transcripts, respectively, without cross-reactivity. The expression of human *FOXG1* from SYN1 and RE1 viruses was detected in neurons in various brain areas, including the cortex, striatum, and hippocampus, consistent with the endogenous *Foxg1* expression domains (Fig. 1B,C). Importantly, exogenous human *FOXG1* expression was absent in *Mki67^+^* proliferating progenitors within the DG, whereas endogenous mouse *Foxg1* remained detectable in these proliferating cells (Fig. 1D-G). These results confirm that SYN1 and RE1 vectors successfully deliver neuron-specific *FOXG1* expression in vivo.

### Injection of AAV-SYN1 and AAV-RE1 increases FOXG1 expression in *Foxg1*-cHet brains

*Foxg1* conditional heterozygous (*Foxg1*-cHet; *Foxg1^fl/+^*;*NexCre*) mice, which lack one *Foxg1* allele in cortical and hippocampal excitatory neurons (Goebbels et al. 2006; Cargnin et al. 2018), exhibited a marked reduction in FOXG1 protein in these brain regions (Fig. 2A-C). To determine whether viral delivery restores FOXG1 levels, SYN1 or RE1 viruses were injected ICV at P1, and brains were analyzed at P32 by immunohistochemistry (IHC) assays with an antibody recognizing both mouse and human FOXG1. The administration of AAV-SYN1 and AAV-RE1 resulted in a significant increase in the FOXG1 protein in *Foxg1*-cHet brains (Fig. 2A,C). Notably, viral treatment not only restored mean FOXG1 levels, but also increased the proportion of neurons with high FOXG1 expression (Fig. 2C,D), suggesting that postnatal neuronal gene delivery effectively rescues FOXG1 protein expression levels in vivo.

**Figure 2.**
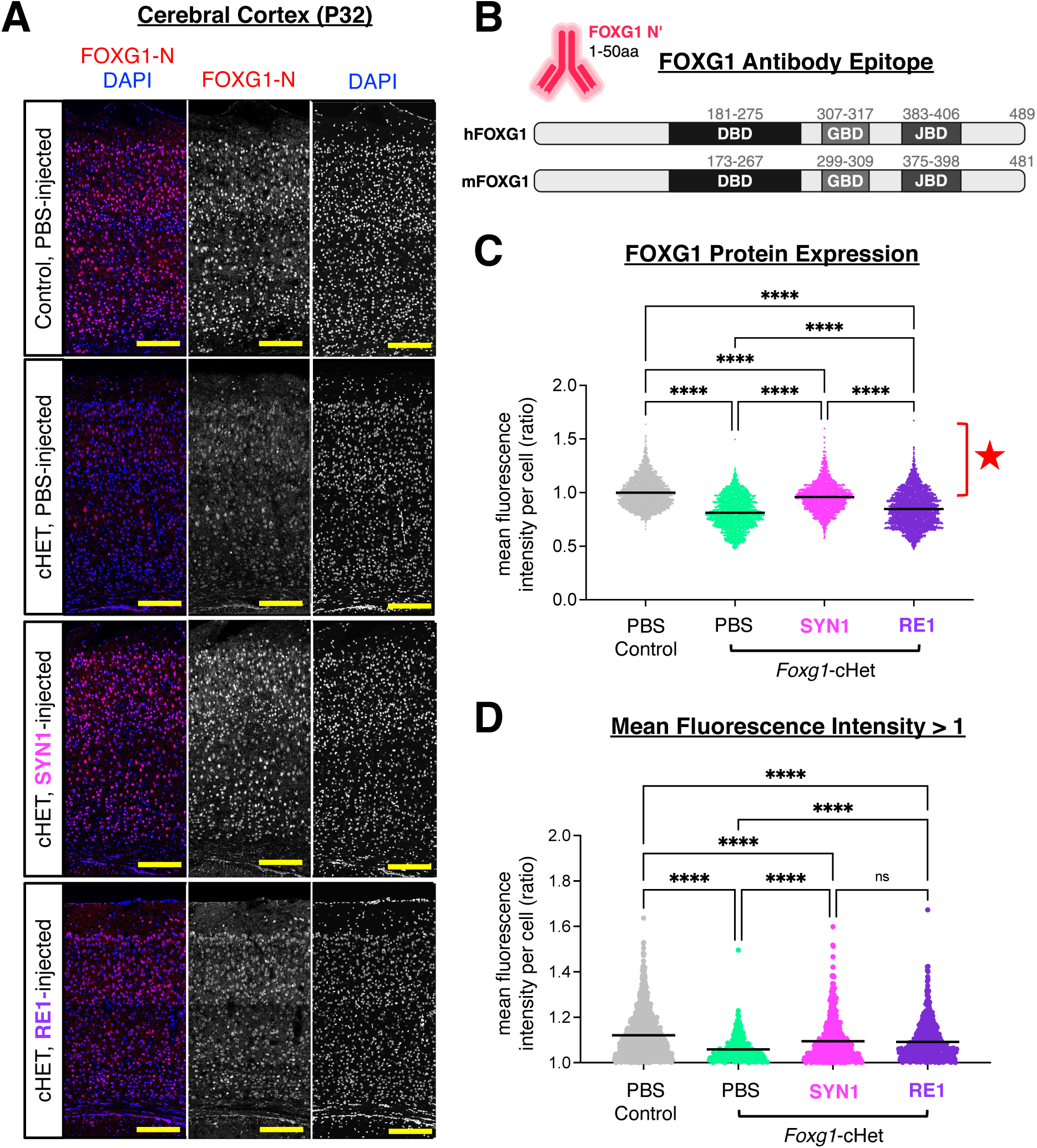
Neonatal AAV-SYN1 or AAV-RE1 treatment restores FOXG1 protein levels in *Foxg1*-cHet mice. (A) Representative images from immunohistochemical analysis of FOXG1 protein levels showing that P1 ICV treatment of SYN1 or RE1 is capable of restoring FOXG1 expression in the *Foxg1*-cHet cerebral cortex at P32. Scale bar 200µm. (B) Schematic representation of FOXG1 antibody binding (Invitrogen PA5-41493), showing that both endogenous and AAV-driven FOXG1 are detected by this analysis. (C-D) Quantification of FOXG1 immunostaining results, graphed as mean fluorescence intensity (MFI) per cell (normalized to control). Bracket and star indicating cells with an MFI greater than 1, which are graphed separately in (D), to show that cells expressing high levels of FOXG1 (which are lost in the *Foxg1*-cHet cortex) are rescued by SYN1 and RE1 treatment. **** p < 0.001; ns, not significant in one-way ANOVA testing. n = 3-4 mice per condition.

### Efficacy of AAV-SYN1 and AAV-RE1 in *Foxg1*-cHet mice

We next tested whether neuronal FOXG1 expression after birth ameliorates brain structural and glial phenotypes in *Foxg1*-cHet mice. Untreated *Foxg1*-cHet mice displayed DG malformations, including reduced length and abnormal apex angles, and oligodendroglial lineage defects characterized by OPC accumulation and reduced myelination (Jeon et al. 2024) (Fig. 3C-E). Neonatal ICV injection of SYN1 or RE1 corrected DG apex length and angles (Fig. 3A–E, I) and normalized oligodendroglial homeostasis by reducing excess OPCs and increasing MBP^+^ myelinated areas in the cortex (Fig. 3F–J). These results demonstrate that neuronal FOXG1 restoration after birth is sufficient to correct both hippocampal architecture and oligodendroglial deficits associated with *Foxg1* haploinsufficiency (Fig. 3J).

**Figure 3.**
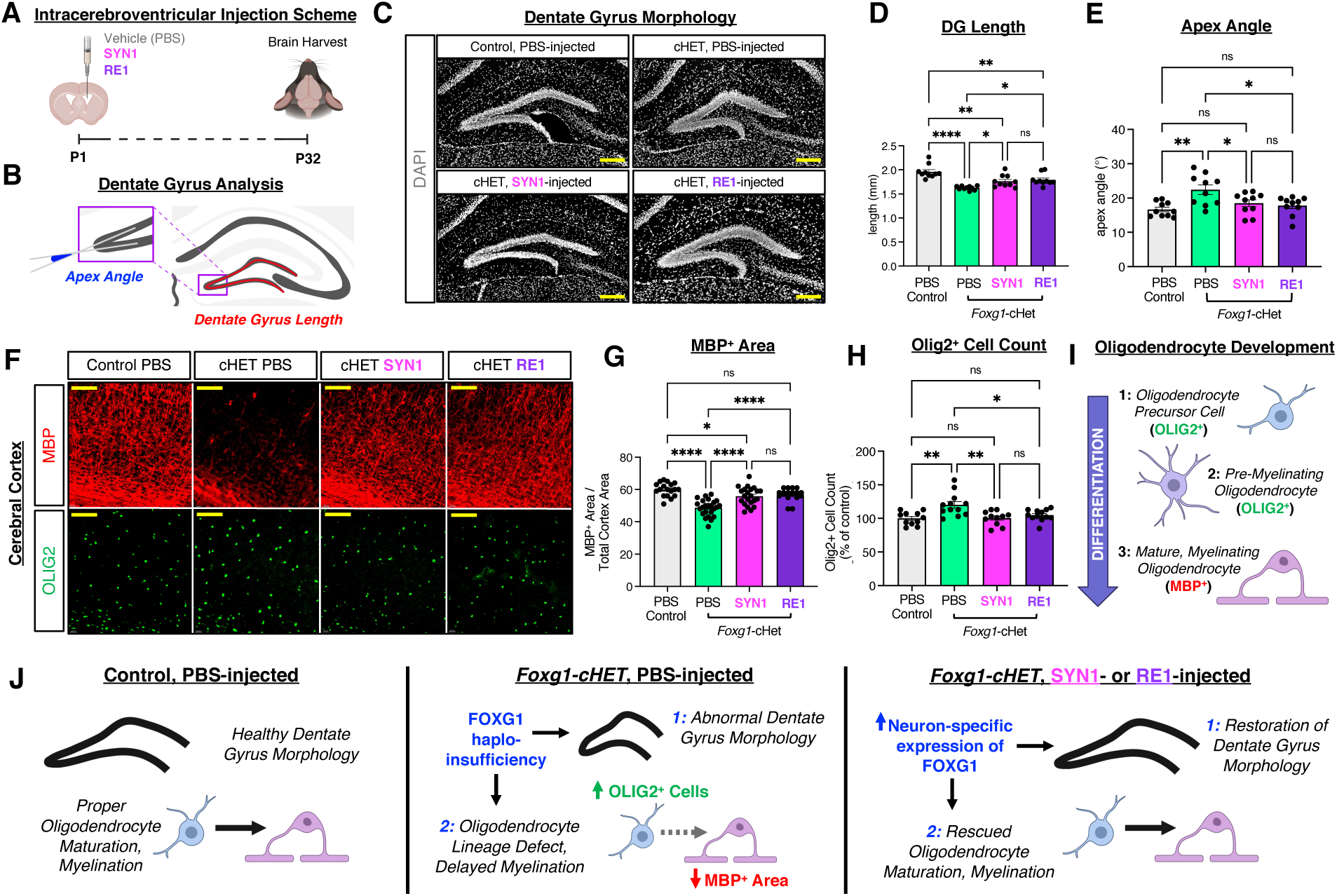
Neonatal AAV-SYN1 or AAV-RE1 treatment restores dentate gyrus morphology and oligodendrocyte lineage defects in *Foxg1*-cHet mice. (A) Schematic representation of neonatal AAV treatment and rescue analysis timeline. (B) Schematic representation of quantification methods used to calculate dentate gyrus (DG) length and apex angle. (C) Representative images showing abnormal DG morphology in the *Foxg1*-cHet is rescued by SYN1 and RE1 treatment. Scale bar 200µm. (D-E) Quantification of DG morphology, showing that reduced DG length and increased apex angle in the *Foxg1*-cHet are rescued by SYN1 and RE1. (F) Representative images showing that aberrant OLIG2^+^ cell production and a corresponding reduction in MBP signal in the *Foxg1*-cHet cortex are rescued by SYN1 and RE1. Scale bar 100µm. (G) Quantification of the MBP+ area within the cerebral cortex (normalized to total cortex area). Area is reduced in the *Foxg1*-cHet and rescued by SYN1 and RE1. (H) Quantification of OLIG2^+^ cell count within the cerebral cortex (displayed as a % of control). OLIG2^+^ cell count is increased in the *Foxg1*-cHet, and restored by SYN1 and RE1. (I) Schematic representation of oligodendrocyte development, showing that oligodendrocyte precursors and pre-myelinating oligodendrocytes are marked by OLIG2, whereas mature oligodendrocytes are marked by MBP. (J) Summary of phenotypic rescue by neonatal SYN1 and RE1 treatment in the *Foxg1*-cHet. * p < 0.05; ** p < 0.01; **** p < 0.001; ns, not significant in one-way ANOVA testing. n = 3-4 mice per condition.

### W300X-Het mice represent a stringent model of severe FOXG1 syndrome

To evaluate the therapeutic efficacy of the SYN1 and RE1 vectors under genetic conditions that faithfully reflect severe FOXG1 syndrome, we employed the mouse line carrying the *W300X* nonsense variant allele (c.924G>A; p.W300X), the precise mouse equivalent of the human *FOXG1* p.W308X (c.923G>A) mutation (Jeon et al. 2025a) (Fig. 4A). The human W308X variant is a recurrent pathogenic nonsense mutation associated with one of the most severe clinical presentations of FOXG1 syndrome, including marked microcephaly, corpus callosum hypogenesis, white-matter volume loss, epilepsy, hypotonia, and profound global developmental delay (Mitter et al. 2018; Lu et al. 2022). Consistent with the severe human phenotype, *W300X* heterozygous mice (hereafter W300X-Het mice) display a broad spectrum of FOXG1 syndrome-related neurodevelopmental abnormalities, including microcephaly, impaired hippocampal architecture, motor deficits, and learning and memory impairments (Jeon et al. 2025a).

**Figure 4.**
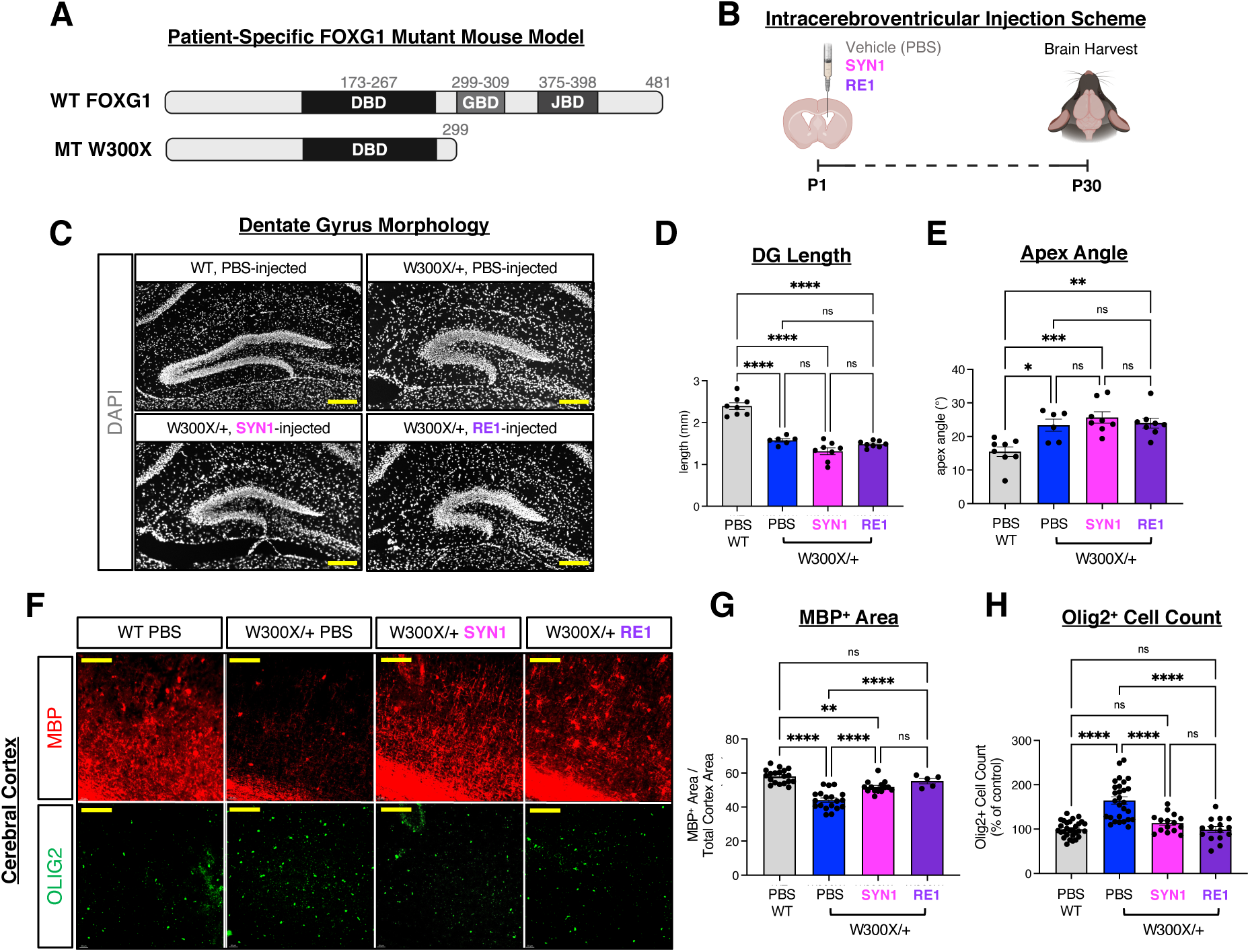
Neonatal AAV-SYN1 or AAV-RE1 treatment restores oligodendrocyte lineage defects, but not dentate gyrus morphology in W300X/+ mice. (A) Schematic representation of the protein structure of WT FOXG1 compared to patient-specific mouse model W300X (equivalent of human W308X). (B) Schematic representation of neonatal AAV treatment and rescue analysis timeline. (C) Representative images showing abnormal DG morphology in W300X/+ mice is not rescued by SYN1 or RE1 treatment. Scale bar 200µm. (D-E) Quantification of DG morphology, showing that reduced DG length and increased apex angle in the W300X/+ are not rescued by SYN1 or RE1. (F) Representative images showing that aberrant OLIG2^+^ cell production and a corresponding reduction in MBP signal in the W300X/+ cortex are rescued by SYN1 and RE1. Scale bar 100µm. (G) Quantification of the MBP^+^ area within the cerebral cortex (normalized to total cortex area). Area is reduced in the W300X/+ and rescued by SYN1 and RE1. (H) Quantification of OLIG2^+^ cell count within the cerebral cortex (displayed as a % of control). OLIG2^+^ cell count is increased in the W300X/+, and restored by SYN1 and RE1. * p < 0.05; ** p < 0.01; *** p < 0.001; **** p < 0.001; ns, not significant in one-way ANOVA testing. n = 3-4 mice per condition.

Importantly, W300X-Het mice express a truncated FOXG1 protein originating from the mutant allele, an aberrant fragment that contributes directly to pathogenesis (Jeon et al. 2025a) (Fig. 4A). Thus, disease mechanisms in W300X-Het mice arise from combined pathogenic processes: loss-of-function (LOF) due to reduced FOXG1 expression, and gain-of-function (GOF) caused by the accumulation of truncated FOXG1 protein (Jeon et al. 2025a). This combination of LOF and GOF mechanisms places the W308X mutation at the severe end of the FOXG1 syndrome spectrum, making the W300X-Het line a stringent, clinically relevant model for assessing the therapeutic potential of AAV-mediated FOXG1 gene replacement.

### AAV-SYN1 and AAV-RE1 exert beneficial effects on both myelination and neuronal phenotypes in W300X-Het mice

To test whether postnatal neuron-restricted FOXG1 expression has efficacy in the FOXG1 syndrome model with severe pathogenesis, we delivered SYN1 and RE1 to W300X-Het mouse brains at P1 via ICV injection. Remarkably, within 30 days after virus delivery, both SYN1 and RE1 suppressed aberrant OPC accumulation and significantly restored myelination (Fig. 4B, F-H). These findings indicate that neuronal FOXG1 expression is sufficient to normalize oligodendrocyte lineage progression and myelination even in this severe genetic context, pointing to non-cell-autonomous contributions of FOXG1 to proper myelination.

To test the efficacy of neuronal FOXG1 expression in alleviating neuronal deficits, we monitored the thickness of cortical layer 6, which contains deep-layer excitatory neurons. W300X-Het cortex exhibited a thinner cortical layer 6 (Supplementary Fig. 2) than the WT control cortex, a hallmark defect previously observed in FOXG1 haploinsufficiency models (Jeon et al. 2025b). Intriguingly, triggering FOXG1 expression in neurons via SYN1 injection at a neonatal stage restored layer 6 thickness in W300X-Het mice (Supplementary Fig. 1), demonstrating that neuronally targeted FOXG1 replacement can correct neuronal deficits despite the presence of a pathogenic truncated FOXG1 protein.

30 days of treatment were insufficient to restore DG structural abnormalities (Fig. 4C-E), likely reflecting the intrinsic severity of the W300X mutation, which impairs DG morphogenesis at early developmental stages and produces a persistent toxic FOXG1 fragment (Jeon et al. 2025a). Nonetheless, the early neuronal and glial improvements suggest that the therapeutic mechanisms have been initiated, but may require a longer window to fully manifest in this severe context.

### Neuronal FOXG1 replacement produces progressive and durable benefits with extended treatment

To test whether a longer post-treatment interval reveals additional rescue effects, we analyzed W300X-Het brains four months after neonatal AAV injection (Fig. 5A). Notably, myelination and DG morphological deficits persisted in adult W300X-Het mice (Fig. 5B-G), indicating that these abnormalities do not self-correct during brain maturation. Both SYN1 and RE1 continued to suppress the excess accumulation of OPCs and enhance cortical myelination (Fig. 5E-G), demonstrating durable long-term efficacy in normalizing oligodendroglial lineage imbalance and differentiation.

**Figure 5.**
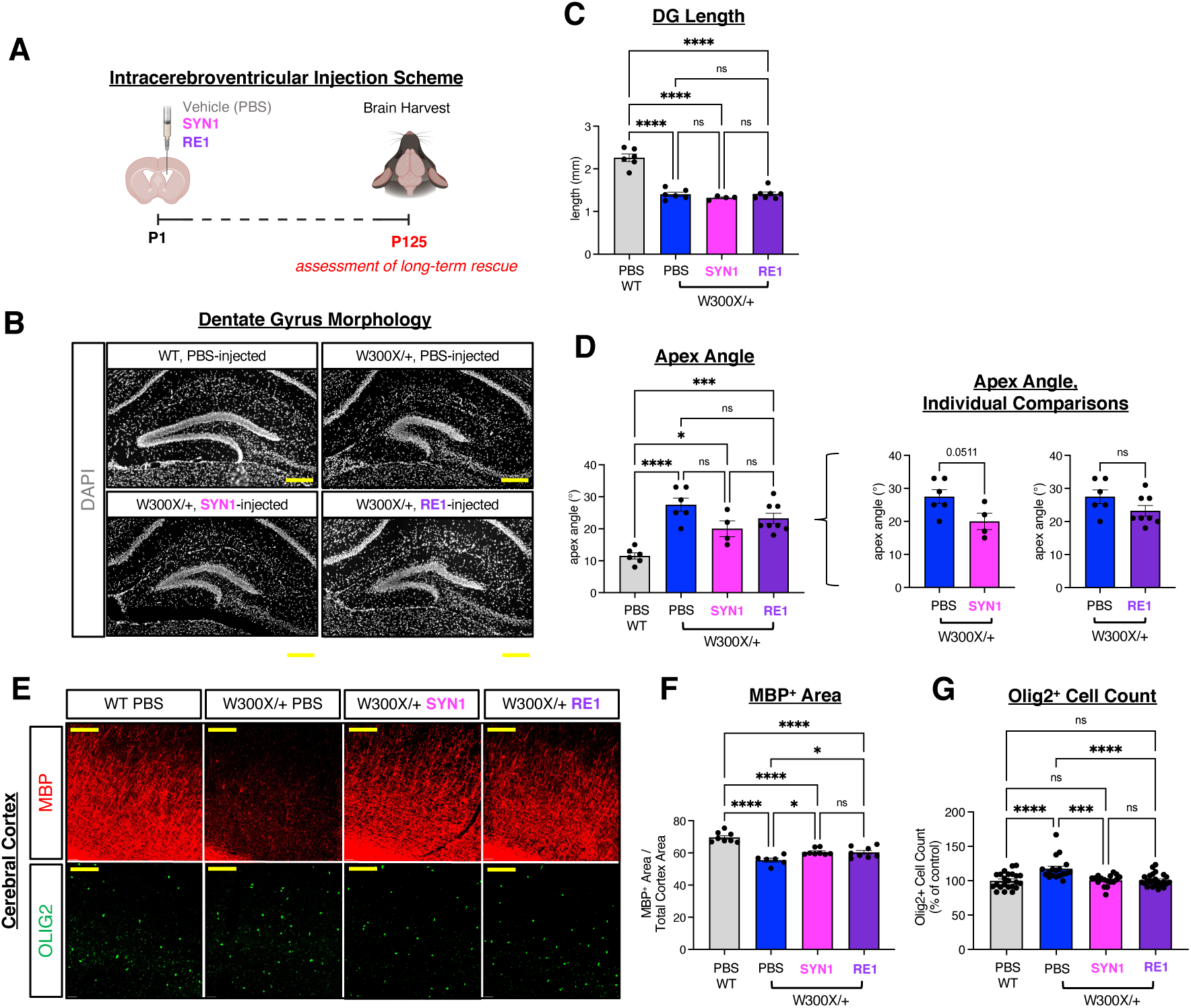
Restoration of oligodendrocyte lineage defects in the W300X/+ by neonatal AAV-SYN1 or AAV-RE1 treatment persists into adulthood. (A) Schematic representation of neonatal AAV treatment and rescue analysis timeline. (B) Representative images showing abnormal DG morphology in W300X/+ mice is not rescued by SYN1 or RE1 treatment. Scale bar 200µm. (C-D) Quantification of DG morphology, showing that reduced DG length and increased apex angle in the W300X/+ are not rescued by SYN1 or RE1. (E) Representative images showing that aberrant OLIG2^+^ cell production and a corresponding reduction in MBP signal in the W300X/+ cortex are rescued by SYN1 and RE1. Interestingly, visible myelination defects in the W300X/+ cortex (marked by MBP) persist into adulthood. Scale bar 100µm. (F) Quantification of the MBP^+^ area within the cerebral cortex (normalized to total cortex area). Area is reduced in the W300X/+ and partially rescued by SYN1 and RE1. (G) Quantification of OLIG2^+^ cell count within the cerebral cortex (displayed as a % of control). OLIG2^+^ cell count is increased in the W300X/+, and restored by SYN1 and RE1. * p < 0.05; *** p < 0.001; **** p < 0.001; ns, not significant in one-way ANOVA testing. Student’s t-tests used for individual comparisons in D. n = 3-4 mice per condition.

Importantly, with this extended therapeutic window, DG apex angle abnormalities began to show partial correction trends (Fig. 5B-D), suggesting that neuronal FOXG1 expression gradually mitigates even complex hippocampal phenotypes over time. These observations are consistent with the hypothesis that the W300X allele’s combined LOF and GOF burden slows the pace of phenotypic reversal and requires prolonged neuronal FOXG1 expression to counteract the sustained effects of the truncated protein.

### Adolescent administration of AAV-CBA viruses restores neuronal and glial phenotypes in W300X-Het mice

To test whether *FOXG1* gene therapy retains efficacy beyond the neonatal period, we used AAV-pCBA-FOXG1, in which the human *FOXG1* expression is driven by the ubiquitously active CBA promoter (hereafter, CBA) (Jeon et al. 2024). We administered the CBA virus via intra-cisterna magna (ICM) injection into W300X-Het mice at P30 (Fig. 6A). Thirty days after administration, CBA-driven ubiquitous FOXG1 expression robustly restored both myelination and DG morphology (Supplementary Fig. 2), indicating that *FOXG1* replacement therapy can reverse neuronal and glial abnormalities even when initiated well after early postnatal development, a stage at which many structural brain defects have already stabilized. These findings suggest that the therapeutic window for *FOXG1* gene replacement is considerably broader than previously anticipated.

**Figure 6.**
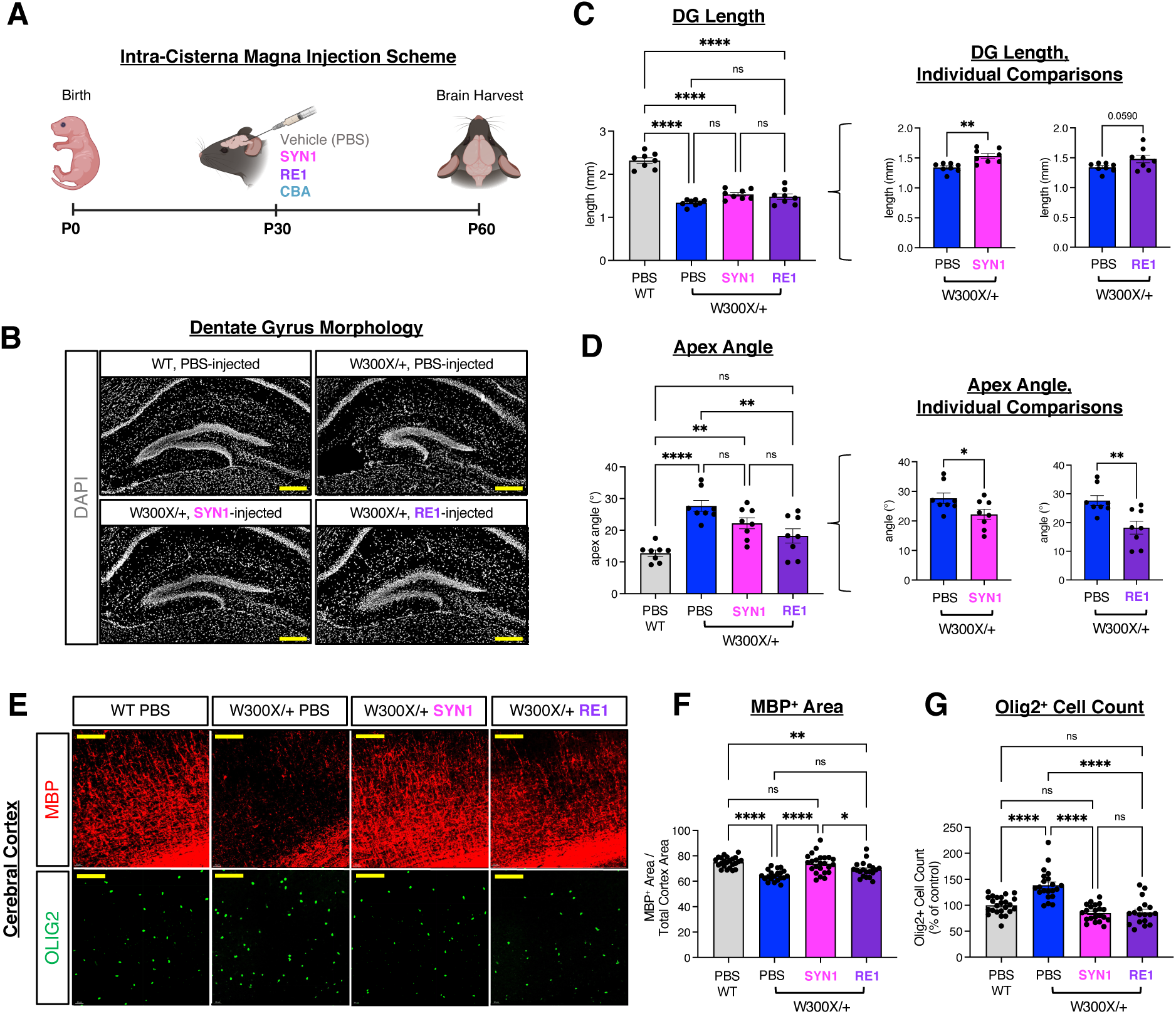
Adolescent treatment of AAV-SYN1 and AAV-RE1 restores oligodendrocyte lineage defects, and partially rescues dentate gyrus morphology in W300X/+ mice. (A) Schematic representation of adolescent AAV treatment and rescue analysis timeline. (B) Representative images showing abnormal DG morphology in W300X/+ mice is partially rescued by SYN1 and RE1 treatment. Scale bar 200µm. (C-D) Quantification of DG morphology, showing that reduced DG length and increased apex angle in the W300X/+ are mildly, but significantly rescued by SYN1 and RE1 treatment. (E) Representative images showing that aberrant OLIG2^+^ cell production and a corresponding reduction in MBP signal in the W300X/+ cortex are rescued by SYN1 and RE1. Scale bar 100µm. (F) Quantification of the MBP^+^ area within the cerebral cortex (normalized to total cortex area). Area is reduced in the W300X/+, fully rescued by SYN1, and partially rescued by RE1. (G) Quantification of OLIG2^+^ cell count within the cerebral cortex (displayed as a % of control). OLIG2^+^ cell count is increased in the W300X/+, and restored by SYN1 and RE1. * p < 0.05; ** p < 0.01; **** p < 0.001; ns, not significant in one-way ANOVA testing. Student’s t-tests used for individual comparisons in C and D. n = 3-4 mice per condition.

### AAV-SYN1 and AAV-RE1 retain therapeutic efficacy in adolescence

We next asked whether neuronally restricted FOXG1 expression remains effective when initiated at P30. W300X-Het mice were injected with SYN1 or RE1 via ICM at P30 and analyzed in adulthood. Adolescent delivery of SYN1 or RE1 significantly reduced OPC overaccumulation and enhanced cortical myelination (Fig. 6E-G). Notably, the magnitude of myelin rescue was comparable to that observed after neonatal administration, underscoring the potency of neuronally expressed FOXG1 in promoting oligodendrocyte differentiation and myelin formation. This is particularly striking given that SYN1 and RE1 drive expression exclusively in neurons, yet rescue myelination deficits, highlighting a strong non-cell-autonomous effect of neuronal FOXG1 restoration on oligodendrocyte lineage maturation.

Notably, DG length and apex angle abnormalities in W300X-Het mice also exhibited significant partial correction by adolescent treatment of SYN1 and RE1 (Fig. 6B-D), demonstrating that hippocampal structural phenotypes remain responsive to neuronal FOXG1 restoration even in late developmental stages. Moreover, adolescent SYN1 administration fully restored the thickness of L1⁺ axonal bundles in the cortical white matter (Supplementary Fig. 3A), correcting an axonal tract deficit that reflects the corpus callosum hypogenesis commonly seen in FOXG1 syndrome. The degree of axonal rescue was complete, suggesting that axonal projection deficits may retain considerable plasticity in the W300X-Het brain, even after adolescence.

Together, our data show that despite the severity of the W300X allele, which combines *Foxg1* haploinsufficiency with the accumulation of a truncated FOXG1 fragment, both SYN1 and RE1 vectors displayed pronounced therapeutic efficacy.

### Neuronal FOXG1 expression is capable of suppressing microglial hyperactivation in W300X-Het brains

A novel and biologically meaningful phenotype observed in W300X-Het mice was the robust increase in microglial activation (Jeon et al. 2025a), as indicated by elevated numbers of IBA1^+^ cells (Supplementary Fig. 3B). Remarkably, adolescent delivery of the neuron-selective SYN1 vector normalized aberrant microglial activation, reducing IBA1^+^ cell numbers to near-WT levels (Supplementary Fig. 3B). This rescue occurred despite FOXG1 expression being restricted to neurons, further emphasizing the non-cell-autonomous mechanisms by which neuronal FOXG1 restoration attenuates glial pathology. These results reveal microglial dysregulation as an additional pathological layer in the W300X-Het model, and establish neuronal FOXG1 expression as a potent suppressor of neuroinflammatory responses. Significantly, our findings broaden the therapeutic window for AAV-FOXG1 treatment, showing that neuronal FOXG1 delivery remains effective in adolescence in rescuing neuronal, myelination, and neuroinflammatory deficits.

## DISCUSSION

Our study identifies neuronal FOXG1 as a central regulator of both neuronal and glial homeostasis and demonstrates that restoring FOXG1 expression selectively in neurons is sufficient to ameliorate a broad spectrum of cellular and structural abnormalities in mouse models of FOXG1 syndrome. By examining both a conditional *Foxg1* loss-of-function model (*Foxg1*-cHet) and W300X-Het mice, which model a severe form of FOXG1 syndrome, we delineate how FOXG1 expression level in neurons shapes neuronal maturation, oligodendrocyte lineage progression, myelination, and microglial activation. Our findings have important implications for understanding FOXG1 biology, the pathophysiology of FOXG1 syndrome, and the design of gene replacement therapies.

### Myelination deficits in FOXG1 syndrome

Hypomyelination is a consistent feature of FOXG1 syndrome patients, observed in MRI studies (Harada et al. 2018; Mitter et al. 2018; Vegas et al. 2018; Pringsheim et al. 2019). Our results provide mechanistic support for the idea that reduced neuronal FOXG1 alone can drive this glial phenotype. This aligns FOXG1 syndrome with a broader class of neurodevelopmental disorders in which primary neuronal dysfunction secondarily impairs myelination (Nguyen et al. 2013; Jin et al. 2017). Since proper myelination is critical for axonal conduction, synchronization of neural networks, and cognitive function (Fields 2015; Saab and Nave 2017), defective neuron-glia interactions would significantly contribute to the neurological symptoms of FOXG1 syndrome, including epilepsy, intellectual disability, and motor impairments.

Although clinical MRI reports in FOXG1 syndrome often describe “delayed myelination” that appears to improve or normalize with age, our findings in both Q84Pfs-Het and W300X-Het mouse models show that hypomyelination continues into adulthood (Jeon et al. 2025a; Jeon et al. 2025b). The apparent discrepancy likely results from differences in measurement sensitivity and developmental context. In patients, most evidence for catch-up myelination is based on qualitative T1/T2 MRI across cross-sectional age groups, with limited true longitudinal or quantitative myelin imaging; subtle but persistent white matter abnormalities may therefore be underestimated. In contrast, histological and molecular analyses in mice reveal sustained reductions in the myelinated area and altered oligodendrocyte lineage progression, indicating a stable biological deficit. Moreover, because the phenotype is observed in both truncation (W300X) and frameshift (Q84Pfs) alleles (Jeon et al. 2025a; Jeon et al. 2025b), the persistence of hypomyelination is best interpreted as a general consequence of FOXG1 haploinsufficiency rather than an allele-specific effect. Importantly, this persistence underscores the therapeutic necessity of AAV-FOXG1 intervention for myelination defects beyond early development.

### W300X as a stringent preclinical model of severe FOXG1 syndrome

The W300X allele represents an especially rigorous system for evaluating therapeutic efficacy because it encompasses both LOF (reduced FOXG1 expression) and GOF (toxic truncated protein) mechanisms (Jeon et al. 2025a). Consistent with this dual burden, W300X-Het mice show more profound neuronal, glial, and microstructural abnormalities than *Foxg1*-cHet mice, underscoring that GOF alleles impose a higher therapeutic bar than simple haploinsufficiency. Despite this high-level challenge, AAV-FOXG1 delivery showed the phenotypic correction in W300X-Het mice, although the pace of phenotypic correction following AAV-FOXG1 delivery is slower in W300X-Het mice than in *Foxg1*-cHet mice. These studies validate AAV-mediated gene-replacement strategies in a broad symptomatic spectrum of FOXG1 syndrome-causing variants.

### Progressive rescue of hippocampal phenotypes in the severe W300X model

An important insight emerging from our analyses is that the restoration of FOXG1 in neurons initiates improvement of glial and neuronal phenotypes in W300X-Het mice, but the structural rescue of DG is delayed. Thirty days after neonatal AAV administration, DG morphology remained substantially impaired, suggesting that DG deficits involving multiple cell types are compounded by the ongoing presence of the truncated FOXG1 protein in W300X-Het brains (Jeon et al. 2025a). However, extending the post-treatment interval to four months revealed trends toward normalization of DG apex angle, indicating that the severe W300X phenotype is partially reversible and that meaningful anatomical recovery may require prolonged FOXG1 expression in this genetic context. This progressive improvement suggests that even severe FOXG1 syndrome alleles retain latent plasticity that can be harnessed by sustained neuronal FOXG1 restoration.

### Neuron-targeted AAV-FOXG1 vectors show potent and durable therapeutic efficacy

Both SYN1 and RE1 vectors, which restrict transgene expression to neurons, demonstrated robust therapeutic activity across models and developmental stages. In W300X-Het mice, neonatal delivery restored myelination, normalized OPC accumulation, restored cortical layer 6 thickness, and showed a trend toward hippocampal architectural correction. Remarkably, adolescent administration (P30) produced improvements comparable to, or in some endpoints exceeding, those achieved by neonatal delivery, including complete restoration of L1⁺ axonal bundle thickness and significant normalization of DG dimensions. The demonstration that neuronally targeted FOXG1 expression remains effective in adolescents greatly expands the therapeutic window and is highly encouraging for clinical translation, as diagnosis typically occurs after birth and often after infancy.

The next critical step is determining whether these anatomical and cellular improvements translate to functional benefits. Given the complexity of human FOXG1 syndrome, spanning motor, cognitive, epileptic, and behavioral domains, functional outcome measures will ultimately be best assessed in the context of a clinical trial, where both clinician-assessed as well as patient/caregiver-reported outcomes and advanced neuroimaging can be integrated with safety and biomarker readouts.

### FOXG1 functions through both cell-autonomous and non-cell-autonomous mechanisms

Our findings highlight the dual nature of FOXG1 action in the brain. Neuronal FOXG1 has clear cell-autonomous roles in cortical and hippocampal development (Tian et al. 2012; Cargnin et al. 2018). Here, we show that FOXG1 haploinsufficiency in neurons (*Foxg1*-cHet mice) is sufficient to trigger glial pathology, manifested by OPC accumulation and impaired myelination. At the same time, selective restoration of FOXG1 in neurons powerfully rescues phenotypes in non-neuronal lineages, most prominently oligodendrocytes and microglia, revealing strong non-cell-autonomous influences on glial biology. In both Foxg1-cHet and W300X-Het mice, neuron-restricted AAV-FOXG1 normalizes OPC numbers and promotes myelination, demonstrating that neuronal FOXG1 is a major upstream regulator of oligodendroglial differentiation and maturation. Our study establishes that neuronal FOXG1 is not only essential for neuronal differentiation and circuit assembly, but also indirectly regulates oligodendrocyte maturation through neuron-glia cross-talk.

FOXG1-deficient neurons may provide insufficient trophic support or generate deleterious by-products that disrupt oligodendrocyte lineage progression. The mechanism may involve neuronal activity-dependent signals, including growth factors, cytokines, and metabolites that shape oligodendrocyte differentiation and myelin sheath formation (Simons and Trajkovic 2006; Thornton and Hughes 2020). In addition, FOXG1 haploinsufficiency may induce metabolic or mitochondrial defects in neurons, as suggested by transcriptomic studies of W300X-Het mice (Jeon et al. 2025a), leading to increased reactive oxygen species (ROS) and stress signaling. Oxidative stress in neurons has been shown to impair oligodendrocyte differentiation (French et al. 2009; Spaas et al. 2021).

Importantly, our data do not exclude additional cell-autonomous roles for FOXG1 in oligodendrocyte lineage cells themselves. Conditional deletion of *Foxg1* specifically in OPCs or oligodendrocytes will be needed to dissect whether direct FOXG1 action in these cells contributes to oligodendrocyte lineage progression or myelin formation.

### Neuroinflammation as a core component of FOXG1 Syndrome disease pathology

A novel and biologically compelling finding is the pronounced microglial hyperactivation in W300X-Het mice (Jeon et al. 2025a), which is normalized by neuron-specific FOXG1 expression. Neuroinflammation is increasingly recognized as a major modifier of neurodevelopmental disease severity (Salter and Stevens 2017; Lukens and Eyo 2022). Our results indicate that loss of FOXG1 drives microglial activation through non-cell-autonomous mechanisms, and this activation is capable of being suppressed by neuron-specific delivery of *FOXG1*. This discovery adds a new dimension to FOXG1 biology and suggests that abnormal neuron-microglia communication may contribute to the symptomatology of FOXG1 syndrome. Therapeutically, the ability of SYN1 and RE1 to reverse microglial dysregulation underscores their capacity to modulate both neuronal and inflammatory aspects of the disease.

In sum, our study reveals that FOXG1 insufficiency perturbs hippocampal formation, myelination, and neuroinflammatory tone through a combination of cell-autonomous and non-cell-autonomous mechanisms. By demonstrating that neuron-restricted AAV-FOXG1 corrects these abnormalities, including in a severe GOF model and even when delivered during adolescence, we provide strong evidence that neuronal gene replacement is a promising and clinically attainable therapeutic strategy for FOXG1 syndrome. These findings refine our understanding of FOXG1 function and establish a framework for future translational efforts to treat this devastating neurodevelopmental disorder.

## MATERIALS AND METHODS

### Animals

All experimental procedures were approved by the Institutional Animal Care and Use Committee at the University at Buffalo under protocol number 201900078 and performed in accordance with the NIH guide for the Care and Use of Laboratory Animals. All mice were housed with a 12-hour dark/light cycle at 25°C and 50% humidity with ad libitum access to food and water. The *Foxg1^flox^*, *Nex-cre*, and W300X mouse lines were described previously (Goebbels et al. 2006; Miyoshi and Fishell 2012; Jeon et al. 2024). All experiments were performed on mice with a C57BL/6 background. *Foxg1*-cHet mice were generated by crossing *Foxg1^flox/flox^* mice with *Nex-cre/+*.

### Viral preparation of AAV-pSYN1-FOXG1 and AAV-RE1-pCBA-FOXG1

A gene cassette containing AAV2_ITR-RE1-RE1-pCBA-SV40 intronA-codon-optimized human FOXG1 coding sequence (CDS)-BGH polyA-AAV2_ITR was designed and commercially synthesized (GeneArt, GmbH, Regensberg, Germany), and hereafter referred to as the pAAV-RE1-pCBA-hFOXG1 plasmid. To replace the CBA promoter with the human Synapsin 1 (hSyn1) promoter, the hSyn1 promoter region was amplified via PCR from the pAAV-hSyn-EGFP vector (Addgene #50465) using Phusion^TM^ Plus DNA Polymerase (F630S; Thermo Fisher Scientific, Waltham, MA, USA). The primers used for amplification were AflII-hSyn1-F (5’-AAACTTAAGAGTGCAAGTGGGTTTTAGGA-3’) and hSyn1-EcoRV-R (5’-AAAGATATCCTGCGCTCTCAGGCAC-3’), which incorporate AflII and EcoRV restriction sites, respectively. Both the pAAV-RE1-pCBA-hFOXG1 plasmid and the hSyn1 PCR product were digested with AflII and EcoRV restriction enzymes (New England Biolabs, Beverly, MA, USA). The digested hSyn1 insert was subsequently ligated into the pAAV-RE1-pCBA-hFOXG1 backbone, replacing the CBA promoter to generate the pAAV-hSyn1-hFOXG1 plasmid (Fig 1A). Recombinant single-stranded AAV serotype 9 (ssAAV9) viruses (ssAAV9-hSyn1-hFOXG1 and ssAAV9-RE1-pCBA-hFOXG1) for use in the experiments were produced using the respective plasmids cloned above. Virus production was commissioned to Charles River Laboratories (Wilmington, MA, USA). Schematic representations of both viruses are available in Figure 1A.

### Viral injection

For intracerebroventricular injection (ICV), neonatal pups were injected within 36 hours after birth. ICV injection was performed under cryo-anesthesia using a 10μL Hamilton micro-syringe. Viral vectors or vehicle controls were injected into the lateral ventricle of the right hemisphere. AAV vectors were diluted to a concentration of 1 x 10^13^ vg/mL and injected at a volume of 5μL per mouse. Phosphate-buffered saline (PBS), which served as our vehicle control, was similarly injected at a volume of 5μL per mouse. For intra-cisterna magna injection (ICM), mice were injected at approximately P30. ICM injection was performed under Isoflurane-induced anesthesia using a 10μL Hamilton micro-syringe. Viral vectors or vehicle controls were injected into the cisterna magna. AAV vectors were diluted to a concentration of 1 x 10^13^ vg/mL and injected at a volume of 10μL per mouse. Phosphate-buffered saline (PBS), which served as our vehicle control, was similarly injected at a volume of 10μL per mouse.

### Tissue preparation

Dissected brain samples were fixed in 4% paraformaldehyde diluted in phosphate-buffered saline (PBS) overnight at 4°C. The following day, brain samples were washed with PBS, and then incubated in 30% sucrose at 4°C for 3-5 days until fully equilibrated. Brain samples were then embedded in Tissue Freezing Medium (Electron Microscopy Sciences) and used for cryo-sectioning (CM1950, Leica). Sections were prepared at a thickness of 18μm.

### RNA *in situ* hybridization and image acquisition

Evaluation of human *FOXG1* mRNA expression driven by SYN1 and RE1 injection, endogenous mouse *Foxg1* mRNA expression, and *Mki67* mRNA expression was performed using RNAscope Multiplex Fluorescent Reagent Kit v.2 (Advanced Cell Diagnostics: ACDBio, 323110) and Opal dyes, according to the manufacturer’s instructions. The human *FOXG1* and mouse *Foxg1* probes with no cross-detection were synthesized by ACDBio as previously described (Jeon et al. 2024). The mouse *Mki67* probe (ACDBio, 416771-C3) was used to detect proliferating progenitors. Opal 520 (Akoya Biosciences, FP1487001KT) for mouse *Foxg1*, Opal 570 (Akoya Biosciences, FP1488001KT) for human *FOXG1*, and Opal 690 (Akoya Biosciences, FP1497001KT) for *Mki67* were used at a dilution of 1:800 to develop each channel. Finally, the sections were counterstained with DAPI and mounted with ProLong Gold Antifade Mountant (Invitrogen, P36930) for imaging. The sections were imaged using EVOS M7000 imaging system (Invitrogen). Three mouse brains and at least three sections per brain were used for fluorescent signal analysis.

### Immunohistochemical analysis and image acquisition

Position-matched sections from 3-4 mice per group were stained using standard immunohistochemical techniques. Sections were first incubated with the primary antibody, followed by species-specific secondary antibodies conjugated to fluorophores (Jackson ImmunoResearch). Finally, sections were also counter-stained with DAPI to identify cell nuclei. The primary antibodies used include rabbit anti-FOXG1 (Invitrogen PA5-41493, 1:1000), rat anti-CTIP2 (Abcam AB18465, 1:1000), rat anti-MBP (Millipore MAB386, 1:500), rabbit anti-OLIG2 (Millipore AB15328, 1:1000), and homemade guinea pig anti-OLIG2 (1:1000) (Wang et al. 2020; Wang et al. 2022b). Images were collected and analyzed as previously described (Jeon et al. 2024)

### Statistical analyses

Data were analyzed using GraphPad 10 (Prism, San Diego CA). One-way ANOVAs were performed using genotype as between-subject variables. All bars and error bars represent the mean ± SEM, and significance was set at p < 0.05 (ns, not significant; * p < 0.05; ** p < 0.01; *** p < 0.001; **** p < 0.0001).

### Image preparation

Schematic illustrations were created using BioRender.com.

## ACKNOWLEDGMENTS

We are grateful to Hyeryeong Park, Valli Duvvuru, Muhua Liu, Taeyeon Kim, and Xuefang He for their contributions to this study. Also, we greatly appreciate the efforts and support of the staff and veterinarians of the University at Buffalo LAF. This work was funded by grants from NINDS/NIH (NS111760 and NS100471 to S.-K.L. and NS118748 to S.-K.L. and J.W.L), FOXG1 Research Foundation, and Simons Foundation, as well as the generous startup fund from the University at Buffalo (to S.-K.L. and J.W.L). The authors have no conflicts of interest to disclose.

