## Supplemental Data for "AAV-mediated neuronal expression of FOXG1 restores oligodendrocyte maturation, myelination, and hippocampal structure in mouse models of FOXG1 syndrome"

#### SUPPLEMENTAL FIGURE 1

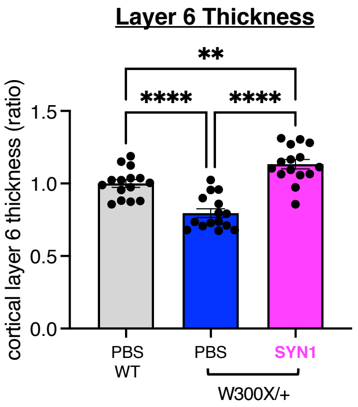


**Supplemental Figure 1. Neonatal AAV-SYN1 treatment restores neuronal layering defects in W300X/+ mice.** (A) Quantification of cortical layer 6 thickness, showing that the significant reduction in layer 6 thickness observed in W300X/+ mice at P30 is rescued by SYN1 treatment administered at P1 via ICV injection. * $p < 0.05$; ** $p < 0.01$; *** $p < 0.001$; **** $p < 0.001$; ns, not significant in one-way ANOVA testing. n = 3-4 mice per condition.

#### SUPPLEMENTAL FIGURE 2

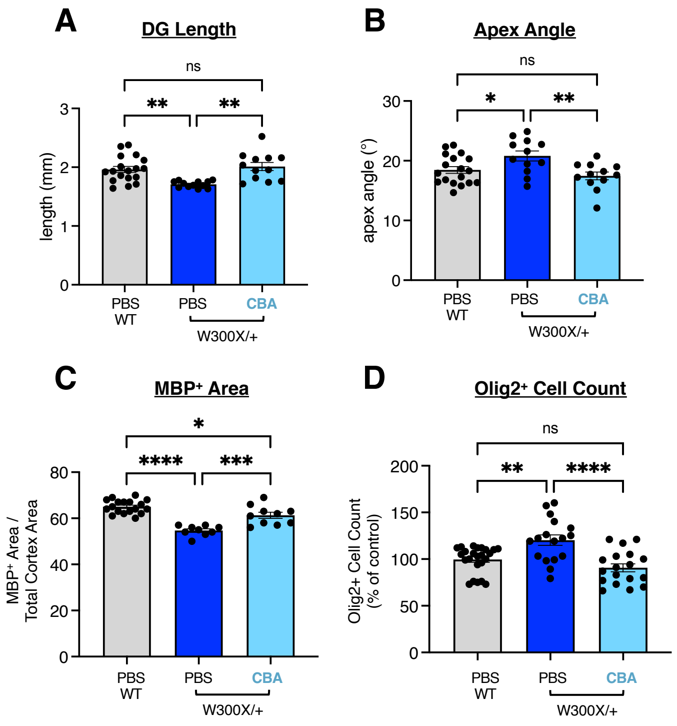


**Supplemental Figure 2. Adolescent treatment of AAV-CBA restores oligodendrocyte lineage defects and dentate gyrus morphology in W300X/+ mice.** (A-B) Quantification of DG morphology, showing that reduced DG length and increased apex angle in the W300X/+ are rescued by CBA treatment at P30 via ICM injection. (C) Quantification of the MBP$^+$ area within the cerebral cortex (normalized to total cortex area). Area is reduced in the W300X/+ and rescued by CBA. (D) Quantification of OLIG2$^+$ cell count within the cerebral cortex (displayed as a % of control). OLIG2$^+$ cell count is increased in the W300X/+, and restored by CBA. * $p < 0.05$; ** $p < 0.01$; *** $p < 0.001$; **** $p < 0.001$; ns, not significant in one-way ANOVA testing. n = 3-4 mice per condition.

### SUPPLEMENTAL FIGURE 3

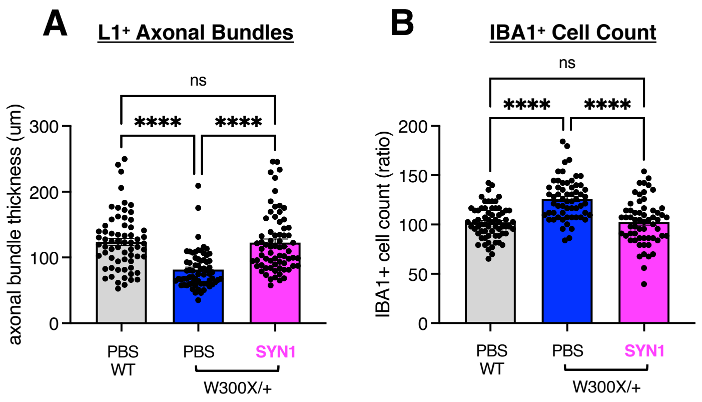


**Supplemental Figure 3. Adolescent treatment of AAV-SYN1 restores axonal bundle thickness and normalized microglial cell numbers in W300X/+ mice.** (A) Quantification of L1$^+$ axonal bundle thickness, showing that reduced axon bundle thickness in the W300X/+ is rescued by SYN1 treatment at P30 via ICM injection (see Figure 6A). (B) Quantification of IBA1$^+$ cell count within the cerebral cortex (displayed as a % of control). IBA1$^+$ cell count is increased in the W300X/+, and restored by SYN1. * $p < 0.05$; ** $p < 0.01$; *** $p < 0.001$; **** $p < 0.001$; ns, not significant in one-way ANOVA testing. n = 3-4 mice per condition.
